# Abundant proteins contribute strongly to noise in the growth rate of optimally growing bacteria

**DOI:** 10.1101/2020.08.28.271668

**Authors:** L.H.J. Krah, R. Hermsen

## Abstract

In bacterial cells, protein expression is a highly stochastic process. At the same time, physiological variables such as the cellular growth rate also fluctuate significantly. A common intuition is that, due to their relatively high noise amplitudes, proteins with a low mean expression level are the most important causes of these fluctuations on a larger, physiological scale. Noise in highly expressed proteins, whose stochastic fluctuations are relatively small, is often ignored. In this work, we challenge this intuition by developing a theory that predicts the contribution of a protein’s expression noise to the noise in the instantaneous, cellular growth rate. Using mathematical analysis, we decomposed the contribution of each protein into two factors: the noise amplitude of the protein, and the sensitivity of the growth rate to fluctuations in that protein’s concentration. Next, we incorporated evolution, which has shaped the mean abundances of growth-related proteins to optimise the growth rate, causing protein abundances, but also cellular sensitivities to be non-random. We show that in cells that grow optimally fast, the growth rate is most sensitive to fluctuations in highly abundant proteins. This causes such proteins to overall contribute strongly to the noise in the growth-rate, despite their low noise levels. The results are confirmed in a stochastic toy model of cellular growth.

---

Stochasticity is inherent to the nature of gene expression (39; 30). Stochastic variation in the copy numbers of proteins is observed even under constant external conditions, and between individual cells in a population of isogenic bacteria. The amplitude of this noise in protein expression can be quantified in terms of the coefficient of variation (CV), defined as the standard deviation divided by the mean. Multiple studies, both theoretical and experimental, consider the scaling of the coefficient of variation squared (CV^2^) with mean expression levels (𝔼 [*X*]) (38; 1). For proteins with a low mean expression, noise is dominated by the intrinsic stochasticity (29; 35) of the chemical reactions involved and CV^2^ scales as 1 / 𝔼 [*X*] (33; 12; 5; 38; 10; 43). For higher mean expression, noise levels decrease and eventually reach a plateau, where fluctuations in gene expression are dominated by extrinsic noise, such as division noise or environmental noise. Because of their larger noise levels, lowly expressed proteins are commonly assumed to be particularly important drivers of fluctuations observed in variables at the cellular level, such as the growth rate. At the same time the effects of the relatively small fluctuations of highly abundant proteins have largely been neglected.

Indeed, noisy gene expression is commonly accepted as the dominant mechanism behind the strong phenotypic divergence that has been observed in populations of genetically identical cells (31). In an exponentially growing population of cells, the growth rate of individual cells is distributed surprisingly broadly (37). Since cellular growth rate is often considered as an important proxy for bacterial fitness, the growth rate –and its fluctuations– have received much attention (16; 18; 15; 32).

In a recent study, correlations between a single protein’s expression level and the cellular growth rate have been explicitly measured for living *E. coli* cells. This showed that fluctuations in the protein’s concentration propagated, via their effect on metabolism, to the growth rate (19). In earlier work, our group proposed a mathematical framework to describe these experimental results. In it, noise in the expression of all proteins in the cell together caused cellular fluxes to fluctuate and, therewith, also propagated to the growth rate (20). We introduced so-called Growth Control Coefficients (GCCs) that quantify, to first order, the sensitivity of the cellular growth rate to changes in the expression of a particular protein species - how much “control” this protein species exerts over the growth rate. Using this theory and the underlying GCCs, we were able to reproduce the characteristics of the observed noise propagation (20).

Another line of research has focused on the remarkable ability of bacteria to tune their protein levels in order to grow, under many external conditions, at a near-optimal rate (3; 42; 7; 23; 34; 13). Optimal gene expression and growth has also been an important and fruitful assumption in countless modelling studies and techniques concerning deterministic growth, including Flux Balance Analysis (27; 6; 14). However, so far this observed optimality has never been taken into account when discussing noise propagation from protein expression to growth rate in single cells.

In this work, we therefore consider systems for which evolution has shaped protein expression to be optimal for fast growth. We then obtain predictions for the contribution of each protein to the noise in the growth rate as a function of its mean expression only. The main result, directly opposing common intuitions, is that proteins with a high mean expression are most important for the noise at the cellular level. The argument is, in short, that a protein’s contribution to noise in the growth rate does not only depend on the protein’s noise level, but also on the sensitivity of the growth rate to that protein’s fluctuations – its GCC. We show that when protein expression levels are optimised for growth, abundant proteins bear more growth control than lowly expressed ones. Consequently, noise in highly expressed protein species affects the growth rate strongly, causing them to contribute strongly to noise in the growth rate despite their low noise levels. Lastly, in a simple stochastic toy model of gene expression and cellular growth, we show these results to be robust for noise amplitudes that resemble those found inside living cells.

## Results

To grow, bacteria need to express a certain set of (metabolic) proteins. Together, these proteins create a metabolic flux used to build cellular components and new proteins. Since the cell’s growth rate is limited by this metabolic flux, noise in the expression levels of the proteins involved propagates through the metabolic network to affect the growth rate (19). For any fixed external environment, we therefore assume the existence of an unknown function *µ* (**X)** that describes the instantaneous rate of cellular growth as a function of the copy numbers of all proteins, **X**. The growth-rate is thus a deterministic function of stochastic variables.

To quantify how fluctuations in a the expression of protein *i* affect growth rate *µ*, we use Growth Control Coefficients (20):

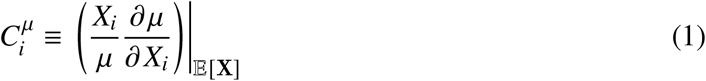

As we will show below, these GCCs offer a way to decompose and analyse the noise in the growth rate in terms of contributions by each of the noisy components, the proteins.

To arrive at a comprehensive and useful noise decomposition, we adhere to two simplifying assumptions. First, noise levels are assumed to be small, so that all protein abundances are close to their means. The growth rate can then be approximated as a linear function of the protein levels. Secondly, fluctuations in all protein species are assumed to be independent. In bacterial cells, this is certainly not the case. However in the case of correlated fluctuations, noise contributions can not be uniquely defined (4; 24). Indeed, when two proteins correlate, and their joint fluctuations affect growth, the attribution of the noise contribution to either protein is arbitrary. Therefore, we here present the simplified case where all protein abundances are uncorrelated so that noise contributions can be uniquely defined and understood intuitively.

Under these assumptions noise in the growth rate, 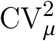, can be approximated as a sum of contributions from all protein species:

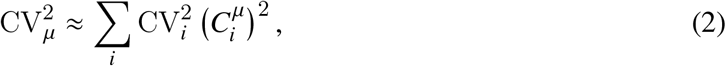

where CV_*i*_ is the coefficient of variation of the copy number of protein species *i* (see Appendix for the derivation).

In this equation, each protein’s contribution consists of two factors. The first factor is no surprise: the proteins’ coefficient of variation quantifies the fluctuations in that particular protein. The second factor is the protein’s GCC, that quantifies how strongly these fluctuations actually affect the cellular growth rate.

### Distribution of Growth Control Coefficients

To further quantify which proteins are important for noise in the growth rate, we need to gain more insight into how growth control is distributed among proteins. This distribution is not arbitrary due to three properties of the GCCs, which are discussed below.

#### Sum Rule

Firstly, the sum of the GCCs equals zero (20):

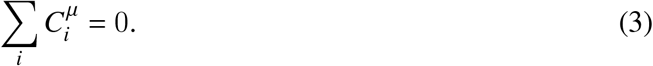

This sum rule originates from the so-called *intensivity* of the growth rate: if all protein copy numbers inside a cell are increased by the same factor, the cellular flux increases, but the mass increases as well, such that the cell’s growth rate (mass increase per mass) stays the same. That the growth rate is to a good approximation an intensive variable has been shown in multiple experiments (19; 37) and is a common modelling assumption (6). Moreover, it is analogous to the assumption that the metabolic flux and cellular mass are *extensive* variables, which is used in Metabolic Control Analysis to derive a similar sum rules for fluxes (17).

#### H-proteins

Secondly, there is a set of proteins, here called *H*-proteins, that are crucial for the cell’s survival, but do not contribute to metabolism or cellular growth. This set *H* includes ‘house-keeping proteins’ participating in, *e*.*g*., stress-response, immunity, and DNA damage repair; in bio-engineering, *H* may also contain engineered pathways. In wild type *E. coli*, the *H*-sector comprises an estimated 25 − 40% of the total protein mass (28).

Even when *H*-sector proteins are not toxic or otherwise harmful to the cell, their control on the growth rate will still be negative. Indeed, while they do not contribute to the metabolic flux, their synthesis does take up resources that otherwise could go to growth-related enzymes. The GCC of such *H*-sector proteins has earlier been calculated (20) to be equal to their mass fraction: 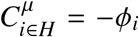. Here we write *ϕ*_*i*_ := 𝔼 [*X*_*i*_] /∑*j* 𝔼[*X*_*j*_] for the proteome mass fractions *ϕ*_*i*_ of each protein species *i* and ignore, for simplicity of notation, that different proteins have different masses. The total *H* sector is denoted as *ϕ*_*H*_.

Together with the sum rule, the presence of the *H*-sector has important consequences for the distribution of GCCs: because some proteins have negative GCCs, others must have a positive GCC.

#### Optimal Growth

Lastly, natural selection tends to favour cells that on average grow faster, which drives the (mean) expression levels of all proteins to be (near)-optimal for growth (42; 7). We here show that this also affects the GCC of the optimised proteins.

To do so, evolution is treated mathematically as a constrained optimisation problem, where the mean growth rate, 𝔼 [*µ*], is optimised under two constraints. First, the cell’s density is kept constant. Second, only a fixed fraction of the proteome (1 − *ϕ*_*H*_) can be allocated towards proteins related to growth. To maximise the growth rate, only the protein abundances inside this fraction can be tuned by evolution while the total size must stay the same.

Using the method of Lagrange multipliers on a linearisation of *µ*, such an evolutionary process can indeed be shown to affect the distribution of growth control between metabolic proteins (for a formal derivation see Appendix). Intuitively, one can reason that in the optimal state all partial derivatives of the growth rate must be equal: if the growth rate would increase more upon increasing 𝔼 [*X*_*i*_] than upon increasing 𝔼 [*X*_*j*_], increasing the expression of *i* at the expense of *j* would increase the growth rate, and hence the growth rate would not be optimal. In terms of the GCCs, this observation translates to:

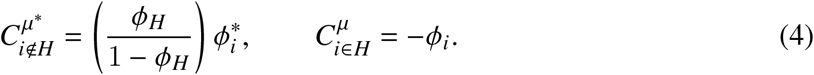

Here, the asterisks indicate that the equality is only valid under optimality.

For cells optimised for growth, Eq. 4 reveals two important properties. First, growth control is shared between all metabolic proteins: there is no single growth-limiting protein. Secondly, and most importantly, enzymes with a higher mean expression level have a proportionally larger control on the growth rate.

### Combining all factors

The use of these insights regarding the distribution of the GCCs in cells growing optimally, combined with the experimentally observed scaling of coefficients of variation of protein levels, results in a prediction for the contribution of each protein to the noise in the growth rate. Moreover, we can analyse how this contribution scales with 𝔼 [*X*], the mean expression level of that particular protein. We write 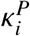 for the predicted noise contribution of protein species *i*, which is defined as the protein’s relative contribution in Eq. 2:

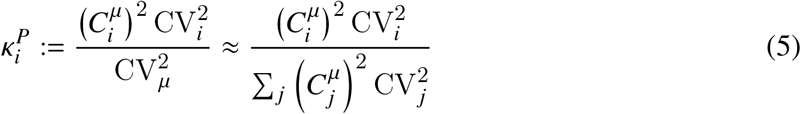

Inspired by experimental data (38; 43), the intrinsic noise component is assumed to be inversely proportional to mean abundance:

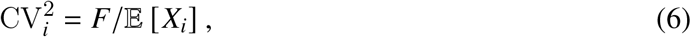

with a fixed Fano Factor *F*. Ignoring the noise plateau caused by extrinsic noise sources, Eq. 6 then sets the noise levels of all protein species. Note that for the highly expressed proteins, noise levels are thus deliberately underestimated, resulting in a conservative estimate for their contribution to noise in the growth rate. To now analyse noise propagation in optimally growing cells, Eqs. 4 and 6 are inserted in Eq. 5. This results in:

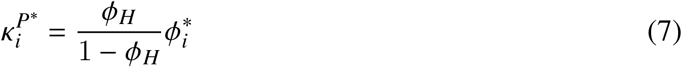

This equation is the pivotal finding of this study. It predicts that, in cells whose expression levels are optimised for growth, proteins with a high mean expression contribute most strongly to fluctuations in the growth rate.

### Stochastic Toy Model

We set out to test the above predictions in a toy model. To do so, we defined a highly simplified model of a growing cell with stochastic protein expression levels. To mimic the effects of evolution, we then employed an optimisation algorithm to search for the mean protein expression levels that optimise the mean growth rate of such cells. Next, we characterised the noise propagation in such optimised cells to verify the predictions of Eq. 4 and 7.

The model cell consists of a metabolic pathway of five reactions in a linear pathway that imports an external metabolite (*m*_1_) and converts it to biomass (Fig. 1A). Each reaction is catalysed by a single enzyme species and inhibited by its own product. Additionally, a sixth protein species is expressed that is not metabolically active, representing the *H*-sector. Given the abundances **X** of all proteins, the instantaneous growth rate *µ* is defined as the steady state flux through the pathway divided by the total number of expressed proteins, including the *H*-sector. Note that the growth rate depends non-linearly on all protein abundances. (For more details, see Appendix.)

**Figure 1:**
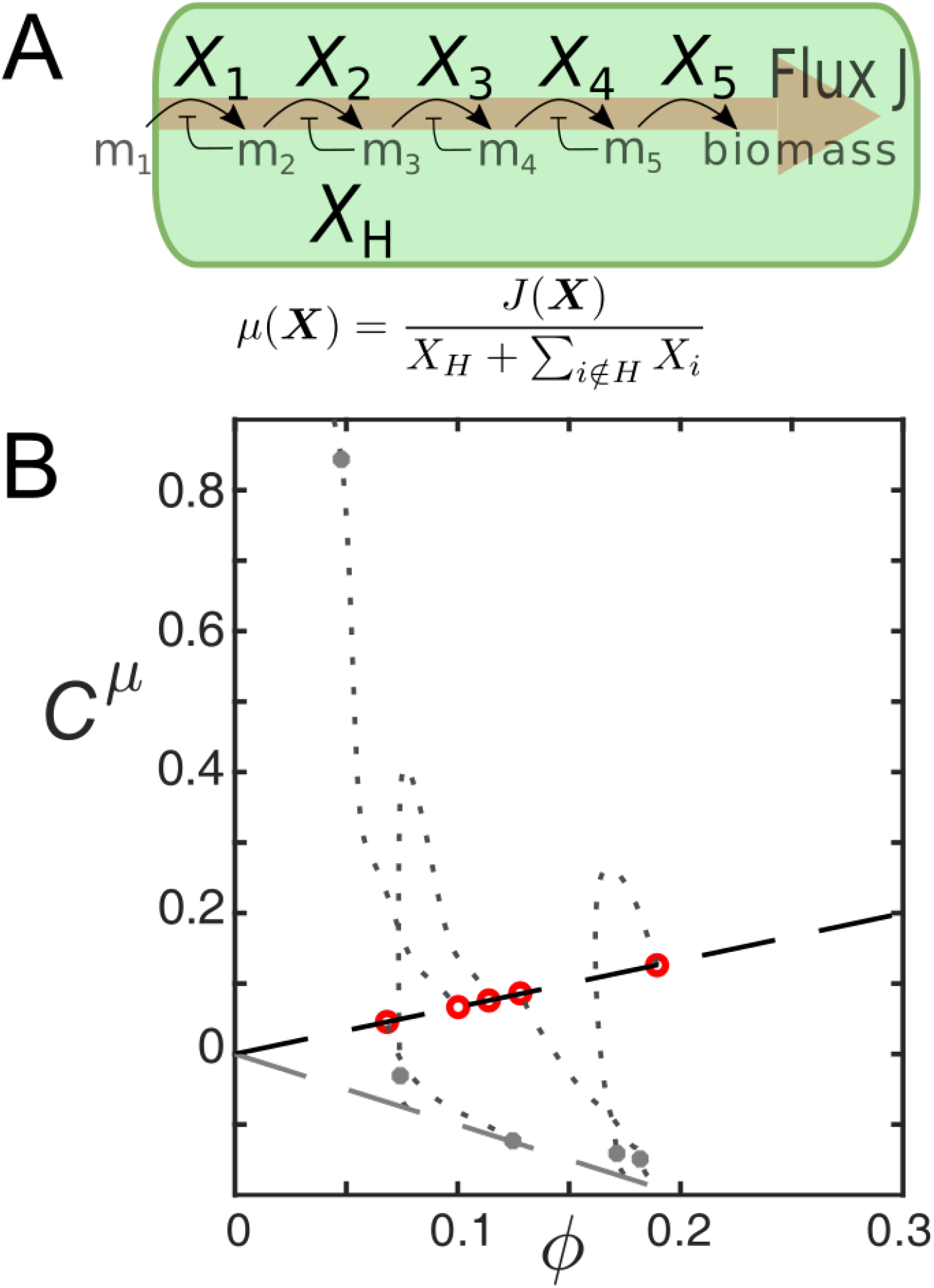
(A) Representation of the stochastic toy model. A cell expresses five metabolic protein species, each catalysing a single reaction in a linear reaction chain that imports and converts a fixed external metabolite, *m*_1_, into biomass. The growth rate is defined as the steady state flux through the network, divided by the total number of expressed proteins, including the *H*-sector protein. (B) Example trajectory during the optimisation of a single kinotype. Grey dotted lines represent the change of the GCCs during the optimisation process. Red point are the values of the GCCs in the optimal genotype, matching the predicted scaling (dashed line). See Tab. 1 for parameters.

The abundances **X** themselves are stochastic: each *X*_*i*_ is distributed according to a Gamma distribution (12) characterised by mean 𝔼 [*X*_i_] and a Fano factor *F*. The Fano factor is chosen the same for all proteins, consistent with Eq. 6, and sets the overall noise amplitude in the cell.

It is useful to distinguish three levels of description of the model cells: their kinotype, genotype, and phenotype. We introduce the ‘kinotype’ as the set of reaction parameters that fully characterise the enzymes in a cell’s metabolic network: kinetic rates, Michaelis–Menten constants and inhibition parameters of the five reactions. We define a cell’s ‘genotype’ as the mean abundances of all protein species. Lastly, a cell’s ‘phenotype’ is given by the vector of the current protein abundances and the co-occurrent growth rate. The phenotype is therefore a stochastic, multidimensional variable whose probability distribution depends on the genotype.

The optimisation algorithm used to mimic evolution searches for the optimal genotype for a given and fixed kinotype. The optimal genotype is the one that generates phenotypes with, on average, the highest growth rate. During an evolutionary trajectory, the genotype is repeatedly subjected to mutations that are subsequently either rejected or accepted. Mutations increase or decrease the mean abundance of one particular protein species, after which the mean expression of all other metabolic proteins is adjusted such that the total cell density remains fixed (∑_*i*_ 𝔼 [*X*_*i*_] = Ω = 10^4^ in all simulations, for more details see Appendix). Mutations are accepted only if they increase the mean growth rate. The mean growth rate of a genotype is determined by sampling many phenotypes generated by that genotype.

After the optimal genotype has been found, the noise contribution of each protein species can be measured by sampling and analysing many phenotypes. Using a method adopted from (4) to measure these noise contributions, they can then be compared with our prediction, *κ*^*P*^ (Eq. 7).

Lastly, the whole process above was repeated for many kinotypes (randomly sampled; see Appendix), resulting in different optimal genotypes.

#### Low-Noise Regime

First we study the system in the low-noise regime (using *F* = 1, Ω = 10^4^), where our results (Eq. 2 and 7) are expected to hold exactly. When noise levels are this low, the mean growth rate 𝔼 [*µ*] is well approximated by the growth rate in the vector of mean abundances, *µ* (𝔼 [**X])**. Instead of using an undirected, slow stochastic optimisation algorithm, the growth rate was therefore optimised with a deterministic gradient-based hill climb algorithm (for details see Appendix).

During each step of the optimisation process, we measured the GCCs of the metabolic proteins to observe how they adjust during optimisation. A representative example of such a trajectory is shown in Fig. 1B (dotted grey lines). Early in the optimisation process, when genotypes are still far from optimal, often one particular protein species is strongly limiting growth (*C*^*µ*^ ≈ 1, indicating that the growth rate could be improved by increasing this protein’s expression level). In contrast, the expression of other metabolic proteins is too high; those proteins have a negative GCC, indicating that most of their expression is a burden to the cell. Eventually, as fitter genotypes are found, growth control becomes shared among all proteins (Fig. 1B, dotted grey lines). When eventually the optimal genotype is found, the predicted positive scaling between a protein’s GCC and its mean abundance is obtained (Fig. 1B, red points, and Eq. 4).

Repeating the same process for multiple kinotypes (*n* = 10) confirms the generality of the positive scaling between *C*^*µ*^ and *ϕ* after optimisation (Fig. 2A, red points). Again, note that in an early stage of the optimisation process the distribution of the GCCs is markedly different (Fig. 2A, grey dots).

**Figure 2:**
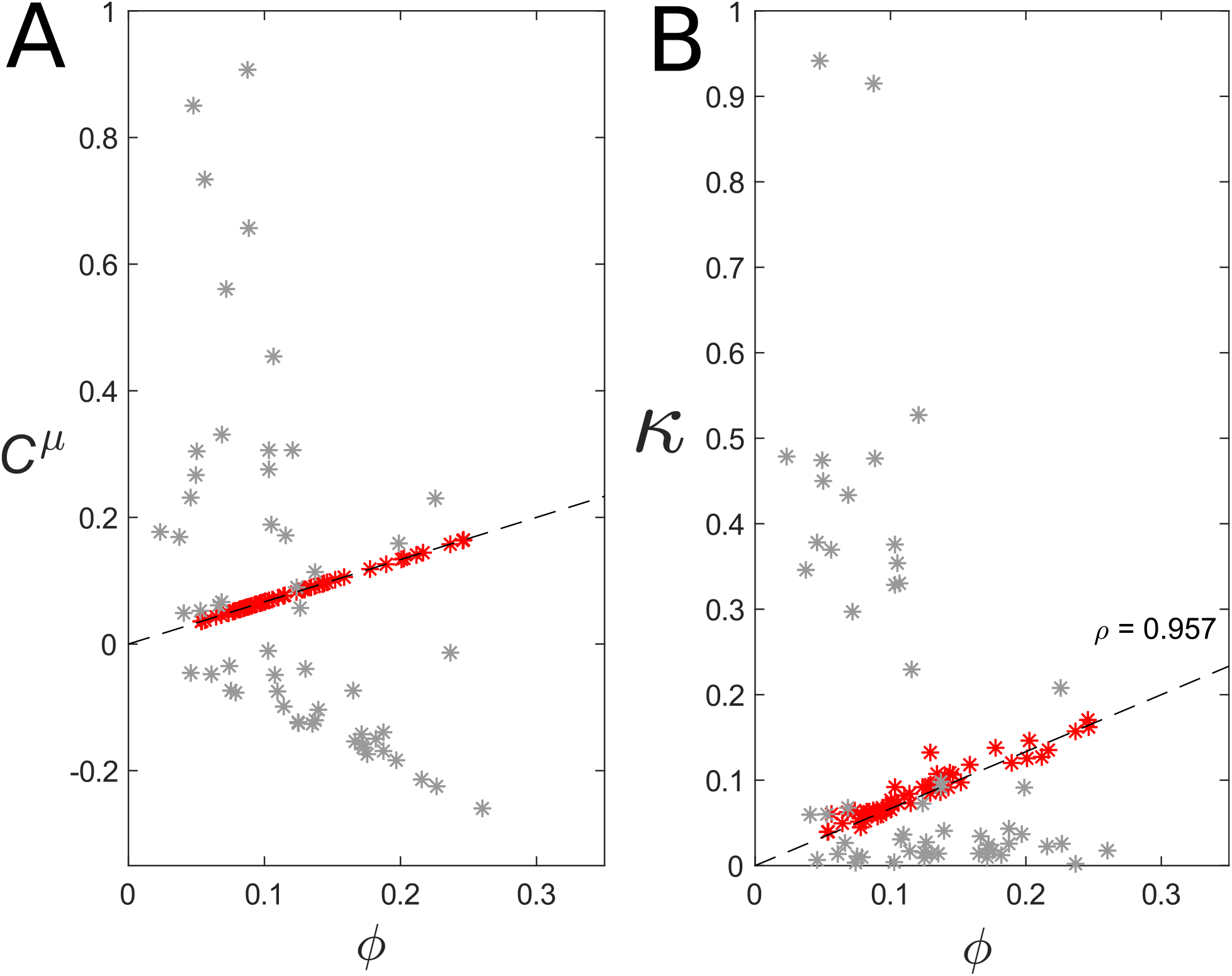
Perfect prediction in the case of linear noise for 10 different kinotypes. (A) Grey dots are GCCs in genotypes after 150 optimisation steps, which are not yet optimal. Red points are the GCCs in the optimal genotypes. Dashed lines are predictions for the values of GCCs for metabolic (positive) or H-proteins (negative) (B) Measured noise contributions compared to predicted noise contributions (dashed line). Red points are measured in the optimal genotype, grey points in the non-optimal genotypes.

Next, we measured for each kinotype the noise contributions of each protein in the optimal genotype, and, for comparison, in a non-optimal genotype. For all kinotypes, the noise contributions in the optimal genotypes neatly follow our prediction (Fig. 2B, red point). In contrast, for non-optimal genotypes, noise contributions are dominated by only a few lowly expressed proteins (Fig. 2B, grey dots). The mean abundance of these proteins is too low, causing both their GCC and their CV to be large, resulting in a large noise contribution.

This analysis clearly highlights the fundamental role of evolution in shaping noise propagation properties: only in evolved cells that grow at an (almost) optimal rate a positive scaling exist between *κ* and *ϕ*.

#### High-Noise Regime

Next we repeat the above analysis for larger noise levels that match those observed in living bacteria (*F* = 10, Ω = 10^4^, resulting in CVs up to 0.2, Fig. S4C). In this regime, the non-linearity of the growth rate becomes important and might affect which genotype is optimal. Below, we therefore distinguish two genotypes for each kinotype: the ‘Low-Noise’ (LN) genotype, which is optimal in the low-noise regime, and the ‘High-Noise’ (HN) genotype’, which evolved in the high-noise regime. To efficiently find the HN genotype of each kinotype, we used the LN genotypes as the starting conditions of the evolutionary algorithm.

For some kinotypes, the resulting HN genotypes indeed differed significantly from the LN ones. This can be understood as follows. When noise levels are low, control over the growth rate is shared between the proteins, and no single protein species is rate-limiting. When noise levels increase while the genotype is kept the same, large fluctuations and non-linear effects can increase the probability that in some phenotypes a single protein species becomes the sole rate-limiting step. Indeed, some genotypes that were optimal in the low-noise regime generated many slow-growing phenotypes when noise levels were high (Fig. S1B). The low growth rates were often caused by a single protein species whose phenotypic expression was too low (Fig. S1A) and therefore became a bottle-neck.

From an allocation point of view, increasing the mean expression of lowly abundant, rate-limiting proteins is cheap: few additional resources are needed to cause a relatively large effect. Indeed, the mean expression of potential bottlenecks increased during evolution, but mainly for lowly abundant protein species (Fig. S1B and Fig. S2).

Unsurprisingly, the distribution of the GCCs is also different in the evolved HN genotype (Fig. 3A, red points). GCCs are by definition linear measures and as soon as the non-linearity of the growth rate become relevant, Eq. 4 is not expected to hold exactly anymore. Interestingly, however, the positive scaling between *ϕ* and *C*^*µ*^ remains, and becomes, if anything, even steeper. Again, this makes sense: when noise levels increase, lowly expressed proteins are, due to their larger CV, more likely to fluctuate down to levels that strongly limit growth (Fig. S2A). Increasing the mean expression of these proteins will reduce their GCC (Fig. S2B-C), while increasing the GCC of the other proteins via the sum rule (Eq. 3). The net result is an increase in the slope in Figure 3A.

**Figure 3:**
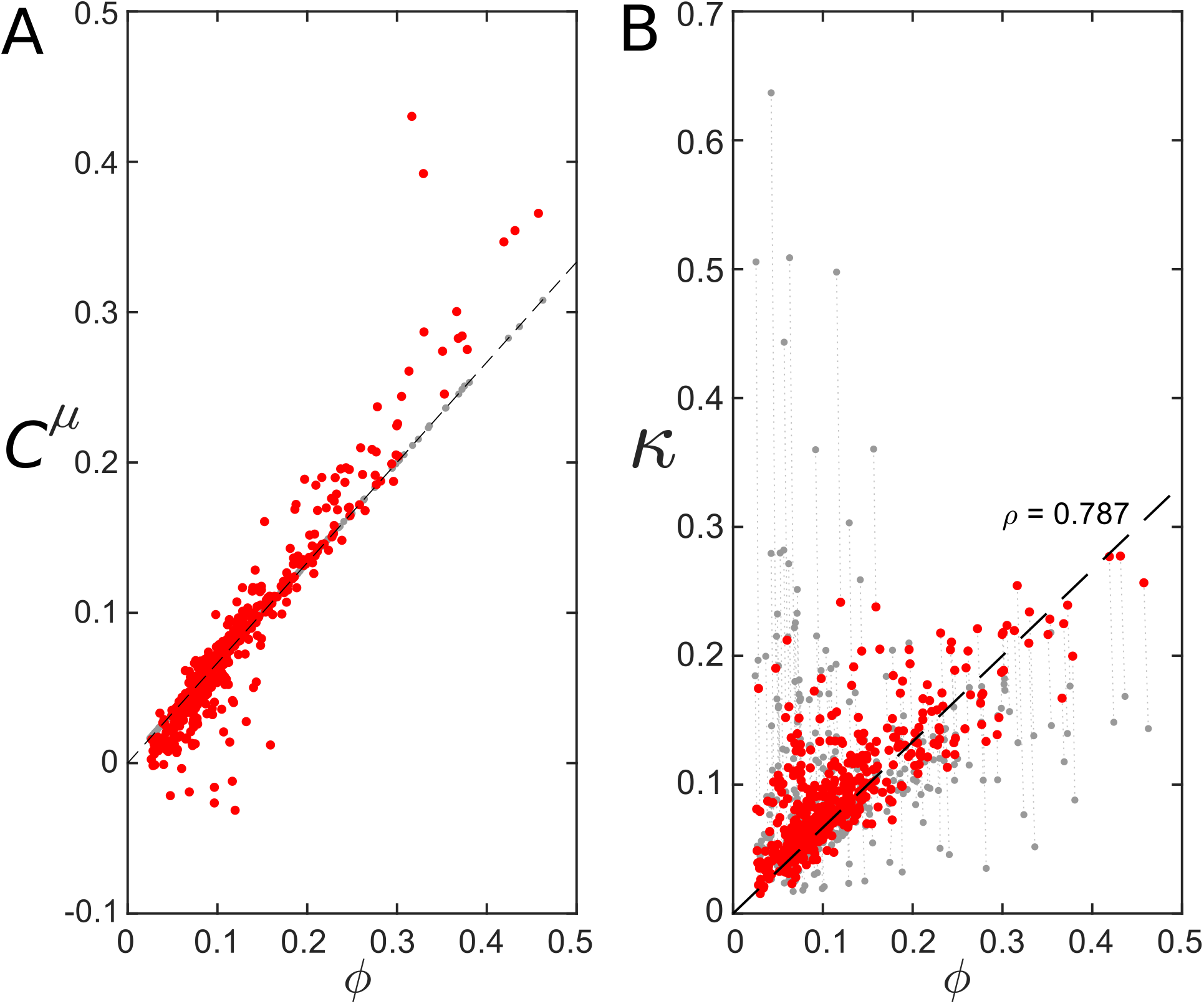
For 100 kinotypes, Low-noise genotypes (grey dots) compared with high-noise genotypes (red dots). Dashed lines are theoretical predictions for metabolic proteins. (A) Growth Control Coefficients. (B) Measured noise contributions.

As growth control and noise propagation are strongly intertwined, much of the same reasoning can be applied to the noise contributions. The positive scaling between *κ* and a protein’s mean expression remained in the high-noise regime (Fig. 3B, red points). At the start of the evolutionary trajectory, when the genotype is still the LN genotype, some lowly abundant protein species are dominant noise contributors (Fig 3B, grey points top-left corner). During evolution those proteins obtained a slightly higher expression, reducing both their GCC and their noise contribution *κ* (Fig. 3B, red points and Fig. S2B-C), and at the same time increasing the contributions of the other proteins species.

## Discussion

In this article we argued that highly expressed proteins play an important role in system-level noise properties. In summary, our argument is that although abundant proteins have low noise levels, their GCCs become large when the cell is optimised for growth. The product of a protein’s CV and GCC -an indicator for the protein’s noise contribution-is predicted to be proportional to the protein’s mean abundance. In a simplistic toy model of cellular growth, we show this proportionality to persist even when noise levels increase and the growth rate is no longer linear.

Although the toy model assumed particular enzyme dynamics, all mathematical predictions were derived without any assumptions concerning the underlying biochemistry. This implies that the strong contribution of highly expressed proteins to the noise in the growth rate is a general property of evolved biological systems.

Furthermore, while this paper only treated a metabolic and an *H*-sector, our analysis can be extended to a cell with additional proteomic sectors, such as the catabolic and ribosomal sectors(21), assuming that their allocation is regulated (26; 44) and they are independently optimised for their specific contribution towards growth (42). On an even smaller metabolic scale, the derived positive scaling between *κ* and *ϕ* will also be present in a single metabolic pathway as long as its protein abundances are optimised, while its total protein abundance is constrained.

The presence of cellular constraints is crucial for our results. In this study, a tight constraint was imposed by assuming that the *H*-sector is completely static. However, a small relaxation of this constraint -*e*.*g*., assuming that the allocation towards the *H*-sector has to be within a certain range-should yield similar results. Also other allocation constraints, *e*.*g*. a maximum density of membrane proteins (36), will result in a similar positive scaling between *κ* and *ϕ*.

We point out that, besides the growth rate, other cellular traits, such as stress response or antibiotic resistance, are important for bacterial fitness as well. Interestingly, noise in these traits can be analysed in the same way as noise in the cellular growth rate. Therefore, the argument suggests that noise in any intensive trait that has been optimised during the bacteria’s evolutionary history should be dominated by highly expressed proteins.

In this paper we focused on ranking the noise contribution of different protein species and ignored other noise sources in the cell. However, other noise sources, *e*.*g*. external fluctuations or division and partition noise, also contribute significantly to variance in system-level variables (25). Therefore, the total variance of the growth rate is underestimated in our simulations, preventing any quantitative predictions for the noise contributions.

Moreover, additional noise sources could cause protein abundances to become correlated (11; 25). Such correlations will remain a problem when discussing noise propagation from the protein-level to cellular growth, because as soon as protein abundances are correlated, noise contributions cannot be uniquely defined. One way to circumvent this problem is to use a fine-grained model description, *e*.*g*. at the level of individual chemical reactions, in which the noise sources are inherently uncorrelated (22; 41), or to adopt a meta-modelling approach (24). That said, the method presented here, by directly relating noise contributions to natural abundances, allows for a more intuitive interpretation of noise contributions.

The results discussed above add to the realisation that global cellular constraints have intricate consequences for the overall physiology of evolved cells, from noisy gene expression (20), to metabolism (36; 14) and growth (44). It highlights the holistic nature of noise propagation via the sum rule for the GCCs (Eq. 3). The sum rule specifically could have important consequences for biotechnology: tinkering with a specific part of the cell affects noise propagation properties of the entire system. For example, most synthetic proteins or pathways do not contribute to growth, but instead create by-products. Such pathways will thus be part of the H-sector, and increase the GCC -and therewith the propagation-of all metabolic proteins.

The stochastic toy model moreover revealed a trade-off between efficient resource allocation and robust metabolism (8). Genotypes that are optimal in the low-noise regime allocate resources efficiently, but lack metabolic robustness at higher noise levels. Cells with different levels of expression noise therefore required different genotypes to grow, on average, the fastest (Fig. 3). Similar observations were also made in a recent experiment in yeast (32). Together, these observations can have consequences for FBA-like techniques, where optimal growth states have so far been calculated mostly deterministically, *i*.*e*., optimising the growth rate in the mean expression levels.

We conclude that noise in gene expression -and its propagation towards the growth rate-needs to be considered when discussing optimal growth, but also vice versa: optimal enzyme expression affects noise propagation properties in such a way that abundant protein species become most relevant for noise on a system level.

## Appendix and Supplementary Figures

### Derivation of Eq. 7

Here we derive equations 2 and (therewith) 7.

#### Linearisation of *µ*

Let the growth rate *µ* be an *intensive* function of the protein copy numbers **X**. Assuming that protein copy numbers are close to some linearisation point 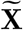, we can write to first order:

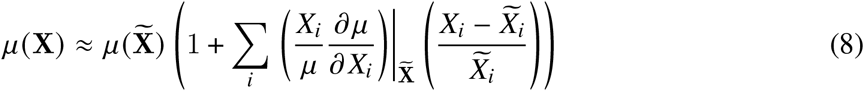

#### Decomposition of CV_*µ*_

When the noise in protein copy numbers is uncorrelated, it is straightforward to calculate CV_*µ*_. First, using the definition of the Growth Control Coefficients of Eq. 1 we can simplify equation 8 by setting 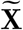 equal to the mean expression levels, 𝔼 [**X**]:

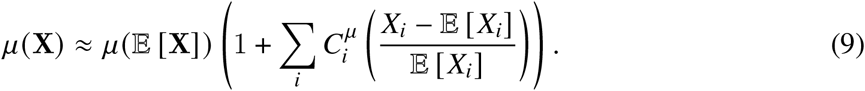

Then, we use the basic properties of the variance, *i*.*e*., for any two uncorrelated stochastic variables *Y*_1_ and *Y*_2_, and scalars, Var { *Y*_1_ +*b* + *Y*_2_} = ^2^Var { *Y*_1_} + Var { *Y*_2_}, to write:

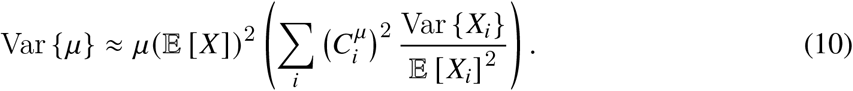

In the regime where Eq. 8 holds, we can also read from Eq. 9 that E *µ* ≈ *µ* (𝔼 [**X])**. This allows us to further simplify Eq. 10:

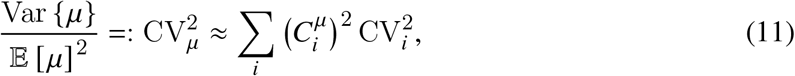

which is Eq. 2 (note that it’s also a special case of (40)).

#### Optimisation

Next, we optimise the *mean* expression levels of the metabolic proteins to achieve the maximal *mean* growth rate. We assume that a fixed number of proteins has to be allocated for other things than growth (the *H*-sector proteins), and constrain the average total cell mass ∑ _*i*_ 𝔼 [*X*_*i*_] to a fixed value Ω. The optimisation can then be written as maximising 𝔼 [*µ*] over 𝔼 [**X** _∉ *H*_], with the constraint ∑_*i*_ 𝔼 [*X*_*i*_] = Ω:

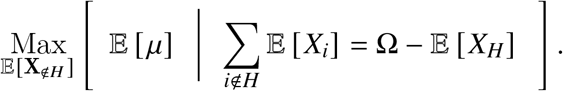

To find the optimal genotype, we use the Lagrange Multiplier method, which in this case takes the form

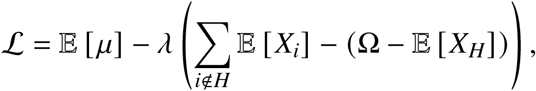

where we need to solve ▽ ℒ = 0. This gives:

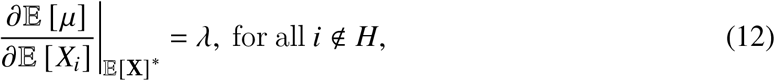

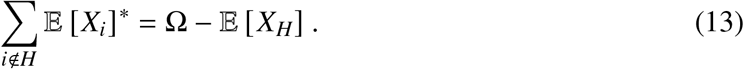

Here the asterisks indicate the optimal value of that variable. Using Eq. 8 we can calculate 𝔼 [*µ*] explicitly:

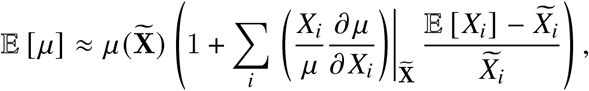

which allows us to calculate the partial derivatives of Eq. 12:

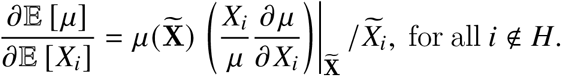

Next, we make a particular choice for the point of linearisation 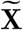 by setting it equal to optimal genotype, 𝔼 [**X**]*. In this vector, the partial derivatives equal *λ* (Eq. 12):

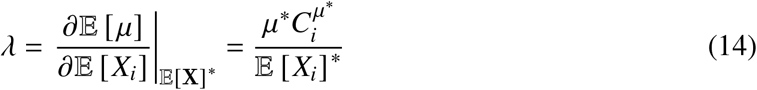

Here, the asterisk denotes that the equation is only valid at the vector with optimal mean expression levels. (Similar arguments were exploited in (2) and (9), although without the inclusion of the *H*-sector and stochastic gene expression.) Lastly, we can use Eq. 13, together with the sum rule for the GCCs (Eq. 3), to calculate, *1* explicitly:

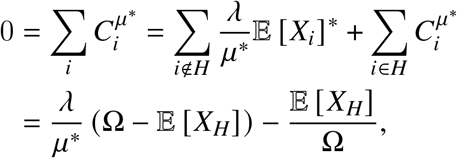

and thus

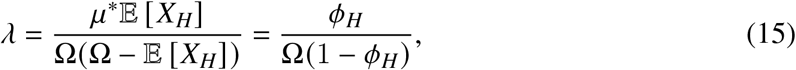

using to the definition of *ϕ*_*i*_ := 𝔼 [*X*_*i*_] */*Ω. Finally, by combining Eq. 14 and 15 we find the Growth Control Coefficient for all proteins that do not belong to the *H*-sector in the optimal growth state:

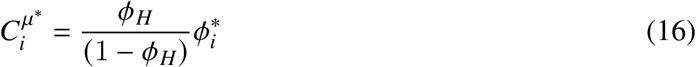

#### Stochastic Simulation

In our toy model, we simulated a linear chain of 5 proteins, where the flux through the first 4 steps is given by:

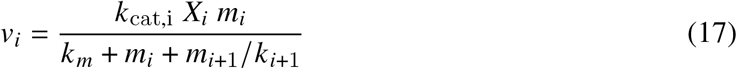

Here, *k*_cat_ is the reaction rate, *k*_*m*_ is the Michaelis-Menten constant, and *k* is the inhibition constant. To define a kinotype, we sample *k*_cat_ and *k*_*m*_ uniformly from the interval 0.1, 6.1 and *k* from 1, 7. The external metabolite, *m*_1_ is set to 10 and kept constant, simulating a fixed environment. The fifth protein in the reaction chain creates biomass and is not inhibited. The initial genotype is constructed by sampling 5 uniform numbers, **x ∈ [**0, 1] ^5^ and setting 𝔼 [**X]** _initial_ = **x***ϕ*_*H*_ */*∑*j x* _*j*_ Ω, with *ϕ*_*H*_ = 0.4 (the proteomic fraction allocated to the H-sector) and Ω = 10^4^, the (mean) total cell size.

Genotypes were mutated in two different ways, depending on the noise amplitude chosen in the simulation. In the low noise regime, the next copy numbers (of all but the *H*-protein) were determined with a gradient-based hill climbing algorithm that uses the current GCCs:

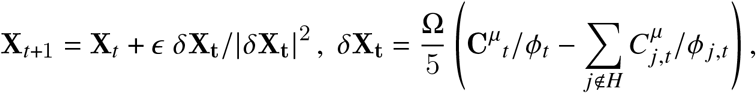

where *ϵ* is small (0.0002). This algorithm changes the genotype in the direction of the steepest growth-rate increase. In the high noise regime, changes in the genotype are stochastic, where one of the metabolic proteins is chosen at random and its mean abundance is changed according to a normal distributed percentage (mean zero, variance 5%), after which the entire genotype is renormalised to enforce a fixed mean cell size of Ω. A mutant genotype is only accepted if it yields a higher mean growth rate. The evolutionary process is terminated when 100 mutants have been declined and repeated 15 times. From these 15 evolutionary trajectories, the genotype with the highest mean growth rate was chosen to be the HN genotype.

A phenotype is then sampled according to six independent Gamma distributions, with means defined by the genotype (𝔼 [**X]** for the metabolic proteins and *ϕ*_*H*_Ω for the *H*-sector) and variance *F*𝔼 [*X*]. Note that although the mean expression levels are always equal to Ω, the total number of proteins expressed in a particular phenotype is not.

To next calculate the growth rate for a specific phenotype, we then integrate the system of ODEs for all metabolites (for *i* ∈ {2, 3, 4, 5}, 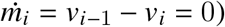 until steady state (10^6^ time steps in Matlab 2019b). Note that the kinetics chosen (Eq. 17) enforce that for all possible phenotypes a steady state exists. In the steady state, where all fluxes are equal, we define *V*_*i*_ = *V*_*i*-1_ = *J*, and:

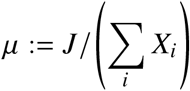

After sampling 2 · 10^4^ phenotypes, we measured noise contributions as follows. First, we divided the distribution for each protein in 100 bins, and calculated the mean growth rate in each bin (sampling extra if less than 100 phenotypes fell in a particular bin). Afterwards, we calculated the weighted variance between these growth rates. This is an approximation of a conceptual decomposition method from Swain *et al*. (4):

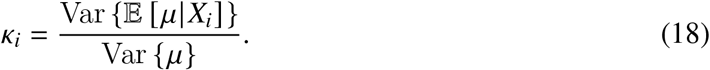

This method is a first order approximation of a full Global Sensitivity Analysis (24). The approximation is only valid if the sum of all contributions is close to unity. This is indeed the case (see Fig. S4A). Moreover, sampling errors in *κ* are small (Fig. S4B). All Matlab codes are available upon request.

**Figure S1:**
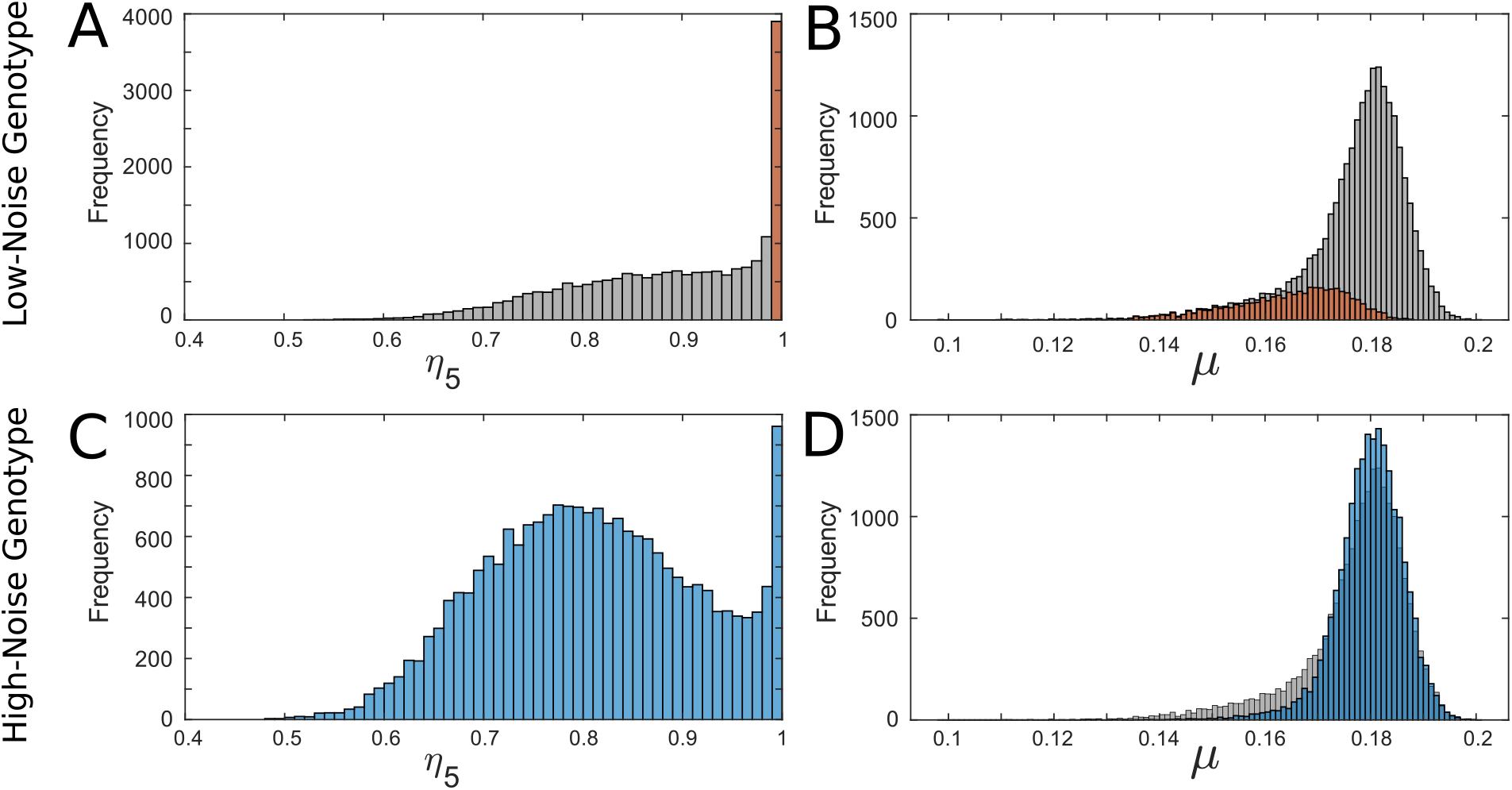
Example of highly skewed distributions in an optimised kinotype. (We picked this kinotype because it most clearly showed the effect of evolution on the distributions.) **(A)** Distribution of the efficiency of the fifth protein 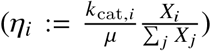. **(B)** Distribution of growth rates for the optimal Low-Noise genotype in the high noise regime. Red areas in (A) and (B) correspond to the same phenotypes. **(C)** The same as (A), but for the evolved High-Noise genotype. **(D)** the same as (B), but for the evolved, High-Noise genotype. Grey distribution is the distribution in the LN-genotype for comparison.

**Figure S2:**
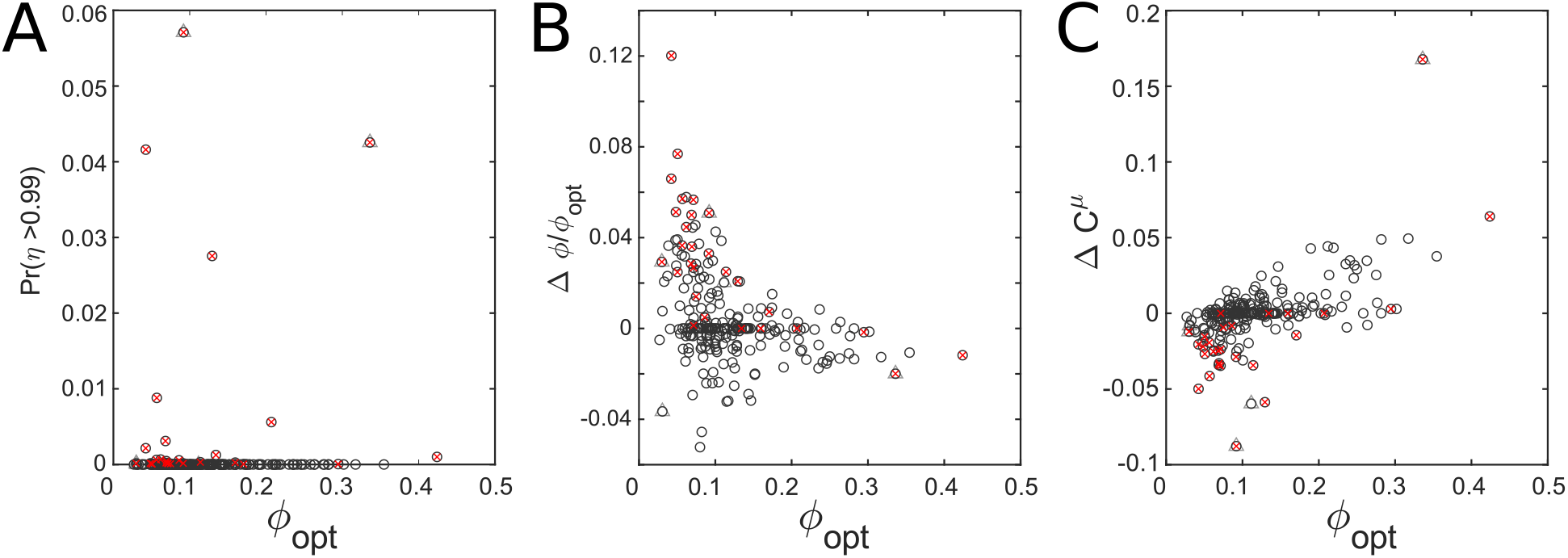
**(A)** Probability a protein’s efficiency is very close to unity, calculated over 2 · 10^4^ phenotypes. Although a higher efficiency on first glance seems good, an efficiency close to unity indicates that this protein might be limiting growth. All proteins for with *p* (*η >* 0.99) *>* 0 are marked with a red cross. Two outliers (triangle) protein species from the same kinotype. **(B)** Changes in genotype when noise levels increase, as a function of the mean *ϕ* in the optimal genotype. Red crosses are those proteins that had a non-zero probability of getting an efficiency close to one in the optimal genotype with high noise. Note that the highly expressed proteins with *p* (*η >* 0.99) *>* 0 are not increased, probably because this required the allocation too much additional resource. **(C)** Changes in mean genotype coincide with a change in *C*^*µ*^.

**Figure S3:**
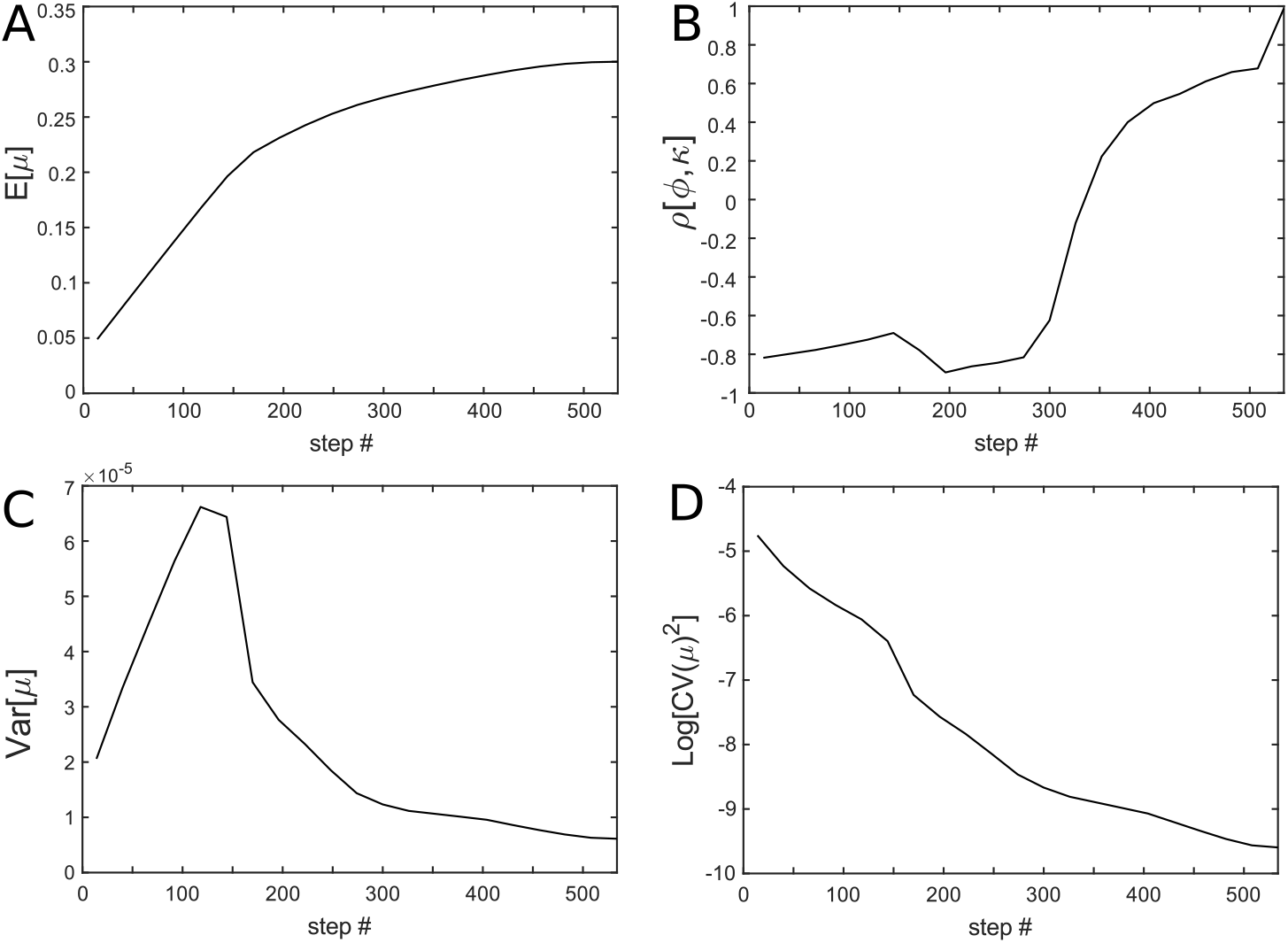
Example of the optimisation algorithm (gradient-based hill climb algorithm). **(A)** Growth rate increases each step. **(B)** Correlation coefficient increases and switches sign during optimisations. **(C)** Variance decreases during most parts of the optimisation process. **(D)** 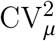 decreases. Parameters of this example kinotype are equal to figure 1B.

**Figure S4:**
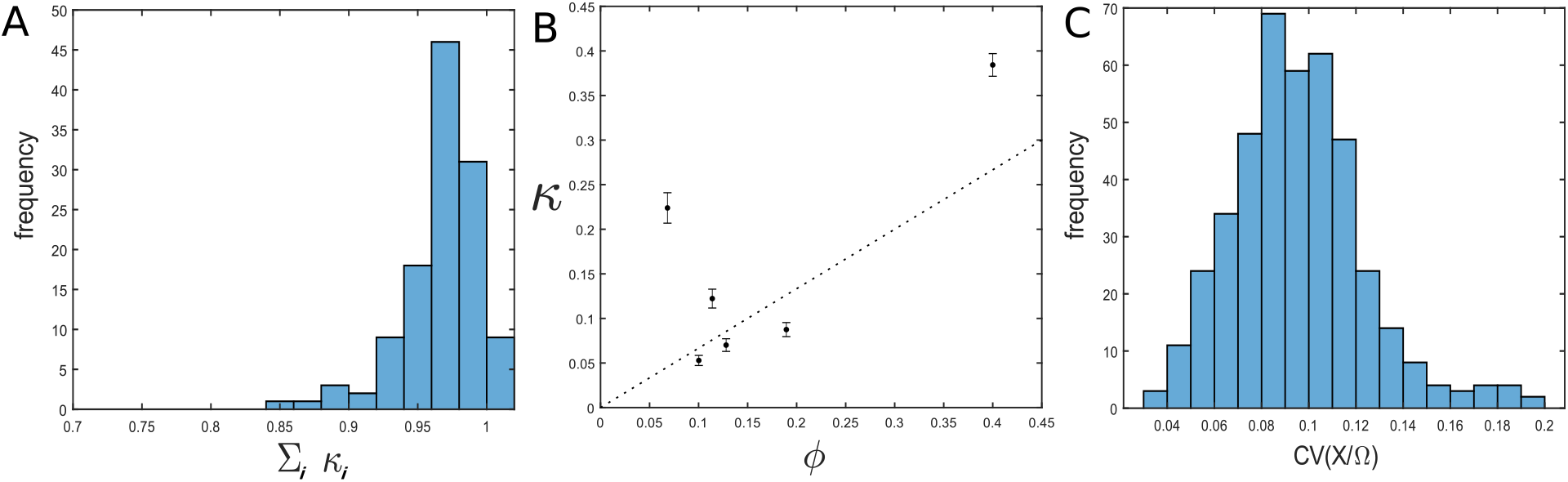
Examination of the noise decomposition method and noise levels. **(A)** The sum of all first order noise contributions is close to unity, indicating that a first order Global Sensitivity Analysis captures the variance contributions well. **(B)** Noise contributions for the example kinotype in the LN genotype in the high noise regime. Error bars indicate 2 sd over 125 repeated sampling of 2 ·10^4^ phenotypes. Sampling errors in *κ* are within reasonable bounds. **(C)** Distribution of CVs as encountered in HN genotypes.

**Table 1:** Kinetic parameters, *k*_cat_ and inhibition constant (*k*_*inhi*_) for the five metabolic protein species in the kinotype used to create figure Fig. 1 and S3. Additionally, the initially sampled (relative) protein abundances are given.

| Protein | $k_{\text{cat}}$ | $k_{\text{inhi}}$ | $\phi_{\text{initial}}$ |
| --- | --- | --- | --- |
| 1 | 3.4048 | 0.2153 | 0.19177 |
| 2 | 4.3489 | 4.3831 | 0.0080544 |
| 3 | 1.8454 | 3.6435 | 0.18137 |
| 4 | 3.1650 | 6.5335 | 0.084103 |
| 5 | 5.4577 | 1.8425 | 0.1347 |

